# *TPCdesign*: A guide to designing informative thermal performance experiments

**DOI:** 10.64898/2026.09.29.755419

**Authors:** Kelsey Lyberger, Lucas A. Nell, Krijn Paaijmans

**Author notes:** equal contribution (Kelsey Lyberger, Lucas A. Nell).

## Abstract

1. Thermal performance curves (TPCs) are widely used to quantify temperature dependence of biological traits, but estimating them requires experiments across multiple temperatures. Because these experiments can be costly, researchers must balance the number and placement of temperature treatments against replication within treatments. Evenly spaced temperatures are a common default, but may allocate effort inefficiently when thermal responses are nonlinear or strongly asymmetric.
2. We developed *TPCdesign*, an R package for planning temperature–trait experiments. *TPCdesign* uses a pseudo-Bayesian (robust) D-optimal experimental design to identify sets of temperatures expected to provide the most information about a TPC, while allowing users to incorporate prior knowledge of thermal limits and curve shape, uncertainty in that knowledge, available numbers of treatments, and practical restrictions on temperature spacing.
3. Using simulation-based worked examples, we show that optimized designs can recover TPCs more accurately than evenly spaced designs, particularly for asymmetric curves and when the number of available temperature treatments is modest. Optimization primarily improves resolution of regions where performance changes rapidly and can substantially improve estimation of upper thermal limits while retaining information across the rest of the curve.
4. *TPCdesign* provides a practical framework for allocating limited experimental resources according to the biological response researchers seek to estimate. Rather than prescribing a universal set of temperatures, it allows experimental designs to reflect existing biological knowledge, uncertainty, and logistical constraints, improving the information gained from costly thermal performance experiments.

## Introduction

Climate change has profound implications for ectothermic organisms, whose physiology and life history traits are tightly linked to ambient temperature. Warming and changing patterns of thermal variability can shift rates of development, survival, reproduction, and other performance related traits, altering population growth, species interactions, and disease transmission in temperature sensitive systems (Krenek et al., 2011; Mordecai et al., 2019; Overgaard et al., 2014). A common way to quantify and predict these temperature-driven changes is through thermal performance curves (TPCs), which describe how the rate of a biological process varies with temperature, often rising to an optimum and then declining rapidly as temperatures approach upper limits (Huey & Kingsolver, 1989; Angilletta, 2009). Thousands of such thermal responses have been characterized for hundreds of traits and hundreds of species (Dell et al., 2013).

Demand for these TPCs has fueled the widespread use of temperature–trait experiments, in which researchers measure physiological or life history traits (e.g., development rate, survival, fecundity) across a range of constant or fluctuating temperatures (e.g., Bernhardt et al., 2018; Paaijmans et al., 2013; MacLean, 2019). These experiments underpin many models forecasting ecological and evolutionary responses to climate change, yet there is considerable variation in how they are designed and implemented (Sinclair et al., 2016). Nonlinear thermal responses imply that ecological conclusions can strongly depend on how well experiments resolve the peak, shoulders, and extremes of the curve, especially under variable temperature regimes (Vasseur et al., 2014). There has also been significant effort to optimize the fit of available experimental data (Régnière et al., 2012) as these experiments can be very costly to run. Recent advances have improved the statistical side of TPC estimation, making it increasingly feasible to fit alternative TPC models and quantify uncertainty in parameters and derived quantities such as thermal optima and limits using nonlinear least squares and bayesian approaches (Padfield et al., 2021; Sorek et al., 2024). However, improved fitting tools do not eliminate a central challenge faced before data collection: deciding how many temperature treatments to include and where to place them while working within practical constraints on experimental capacity, time, organisms, and cost.

Because physiological responses often follow nonlinear trajectories, ad hoc or uniformly spaced temperature treatments may waste resources by oversampling less-informative portions of the performance curve, while failing to capture critical thermal limits or optima. Such under- or oversampling can lead to biased TPC parameter estimates or estimates with very large uncertainties. Despite the clear importance of thoughtful temperature selection, researchers currently lack practical guidelines that account for curve shape, which often varies by trait, or how to incorporate informative priors (Molnár et al., 2017). As a result, researchers often default to convenient, evenly spaced temperature increments across a wide range, inadvertently undermining the power of their experiments.

Here, we address this gap by creating the R package *TPCdesign*, which uses simulation-based evaluation to identify temperature sampling strategies that improve estimation of TPC shape and key derived parameters (i.e., upper and lower thermal limits, optimal temperature) under realistic experimental constraints. We translate these results into practical guidance that increases the efficiency of temperature–trait experiments and the robustness of TPCs and strengthens downstream predictions of ectotherm responses to climate change.

### *TPCdesign* R package

*TPCdesign* is implemented as an R package available on GitHub (https://github.com/lucasnell/TPCdesign) and can be installed using the install_github function from the remotes package. The package requires R version ≥ 4.1.0 and depends on packages acebayes (Overstall et al., 2020), lhs (Carnell, 2026), Rcpp (Eddelbuettel & François, 2011), and RcppArmadillo (Eddelbuettel & Sanderson, 2014). Optional multithreading uses the future.apply (Bengtsson, 2021) package. *TPCdesign* provides functions to calculate performance for TPCs, design temperature treatments, and simulate experimental data. The overall workflow is for the user to specify inputs including priors for thermal limits and optimum, the strength of those priors, and estimated data noise and resource constraints, then the program optimizes candidate temperature designs, and outputs the top design (Figure 1).

**Figure 1.**
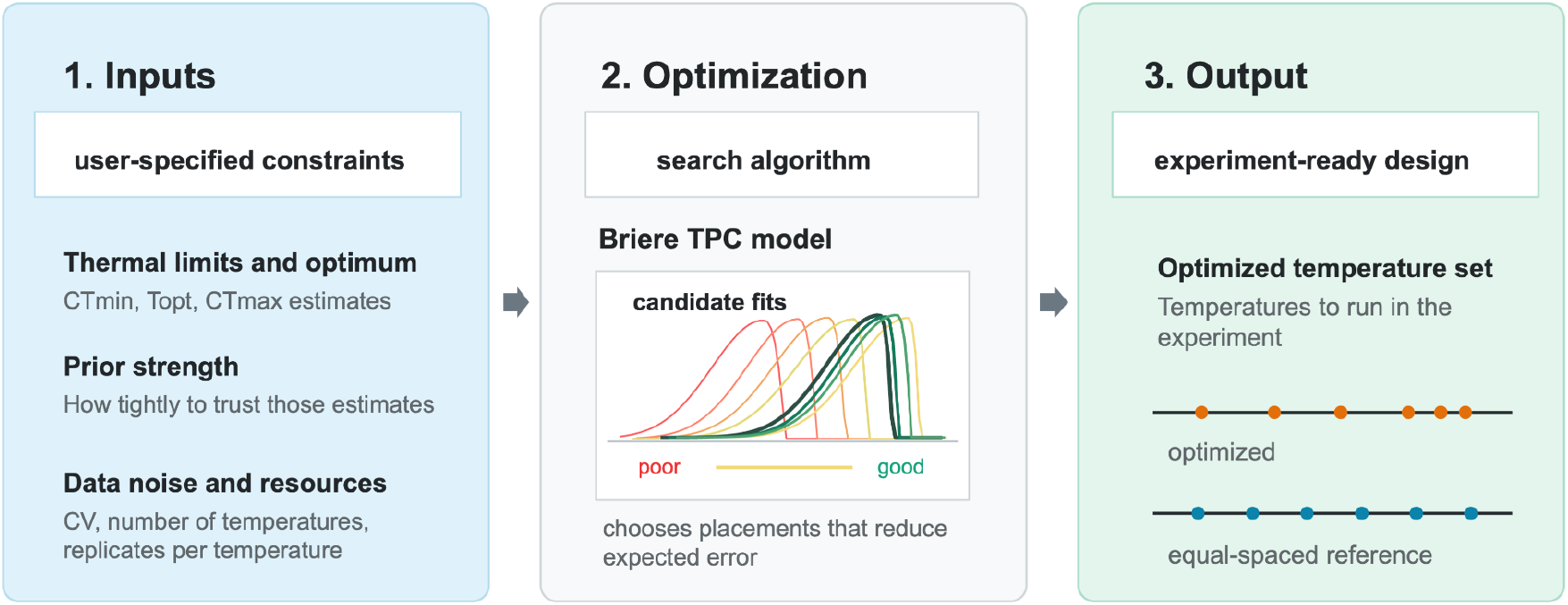
The three steps of the workflow for implementing the *TPCdesign* R package.

Thermal performance was described using the Briere-2 model (Brière et al., 1999). This model uses ecologically interpretable parameters and is commonly used to estimate ecotherm TPCs (Quinn, 2017; Rebaudo & Rabhi, 2018). In our parameterization, performance rises from zero below a critical temperature (*CT*_min_), increases to a peak, and declines relatively sharply to zero above an upper critical temperature (*CT*_max_). Parameters *a* and *b* govern the height and skewness of the curve, respectively. For a given set of parameter values, function briere2_tpc calculates performance across temperatures, and briere2_tpc_Topt calculates the thermal optimum.

To design temperature treatments, *TPCdesign* uses a Bayesian D-optimality framework (Chaloner & Verdinelli, 1995) to choose temperatures that maximize the expected information the resulting data would provide about the four TPC parameters (*CT*_min_, *CT*_max_, *a*, and *b*). For each candidate set of sampling temperatures, we compute the model’s Jacobian with respect to the four parameters at each temperature to form a Fisher information matrix. We then use its log determinant as the D-optimality utility, which is a standard criterion that favors designs that minimize the joint uncertainty in the parameter estimates. A small ridge term (10^−6^) is added to the diagonal of the information matrix before taking the determinant to avoid infinite values when a candidate design is completely uninformative (i.e., all points outside [*CT*_min_, *CT*_max_]).

True TPC parameters are not known in advance, so designs that are optimal for a single parameter set may perform poorly when true values differ. Thus, we extended our approach to a pseudo-Bayesian (robust) design, where we averaged the log-determinants of the Fisher information matrices across a sample of parameter vectors (default 250 draws) drawn from priors reflecting our uncertainty in *CT*_min_, *CT*_max_, and *b*. Priors for *CT*_min_ and *CT*_max_ were uniform distributions centered on user-supplied prior values with user-specified uncertainties as half-widths of the distribution (defaults 2.50 and 1.50 for *CT*_min_ and *CT*_max_, respectively), and the prior for *b* was specified as uniform on the log scale to keep this shape parameter positive (default half-width is 0.26). These defaults were determined using TPCs in Buckley et al., (2021) (Supplement S1). The parameter *a* was held fixed, as it primarily scales the curve and has comparatively little influence on which temperatures are most informative. Averaging the utility over these priors protects the resulting design against misspecification of the user-supplied prior TPC parameters (Chaloner & Verdinelli, 1995).

Rather than optimizing all sampling temperatures simultaneously—a high-dimensional, computationally expensive search—we used the Approximate Coordinate Exchange (ACE) algorithm (Overstall & Woods, 2017), through the R package acebayes (Overstall et al., 2020). ACE updates one temperature at a time conditional on the others, which scales much better with the number of temperatures and is considerably more robust to local optima than jointly optimizing over the full temperature vector at once. To reduce the chance of converging to a local rather than global optimum, we ran ACE from multiple random starting designs (default 7) and retained the design with the highest realized expected utility across starts, implemented as a multi-start, parallelized wrapper around acebayes::ace. Each starting design was generated using a random Latin hypercube sample using lhs::randomLHS (McKay et al., 1979) over the candidate temperature range, which spreads the initial temperatures more evenly across the range than simple uniform-random sampling and thus gives ACE better-diversified starting points.

The user can adjust a number of factors to tune the optimization process to their system, including their certainty in parameter estimates, number of prior draws, and number of starting designs. In addition, design temperatures can have a minimum separation between them (default 2). This prevents the optimizer from placing multiple sampling temperatures arbitrarily close together, which would be statistically informative in theory but impractical or wasteful to replicate in an actual experiment. Lastly, equally spaced filler points (default 1) can be added to hedge against model misspecification and greater-than-expected inaccuracy in model parameters.

### Worked examples

We used simulations to illustrate when optimizing the placement of temperature treatments improves recovery of the underlying TPC. First, we compared optimized and equally spaced designs for curves differing in skew while holding the other parameters constant. For each design, simulated observations were generated at the selected temperatures. To represent experimental error, we simulated replicate observations at each sampled temperature from a gamma distribution parameterized by a constant coefficient of variation (*obs_cv*), such that relative error remained approximately constant across temperatures. The TPC was estimated from these observations. We evaluated overall recovery of the curve (as root mean square error, RMSE, for 101 points from *CT*_min_ to *CT*_max_), as well as estimation of the biologically interpretable quantities *CT*_min_, *T*_opt_, and *CT*_max_. Optimized and equally spaced designs produced similar fits when the true curve was relatively symmetric, but their performance diverged as the curve became more asymmetric (Figure 2). For strongly skewed curves, *TPCdesign* shifted temperature treatments toward regions where performance changed most rapidly rather than distributing them uniformly over the full thermal range. This reduced error across the fitted TPC. The largest improvement occurred in estimation of *CT*_max_, with a smaller improvement in *T*_opt_ and a modest tradeoff in estimation of *CT*_min_ (Figure 2B). Thus, optimization was most consequential when the shape of the TPC created substantial differences in the amount of information provided by different parts of the temperature gradient.

**Figure 2.**
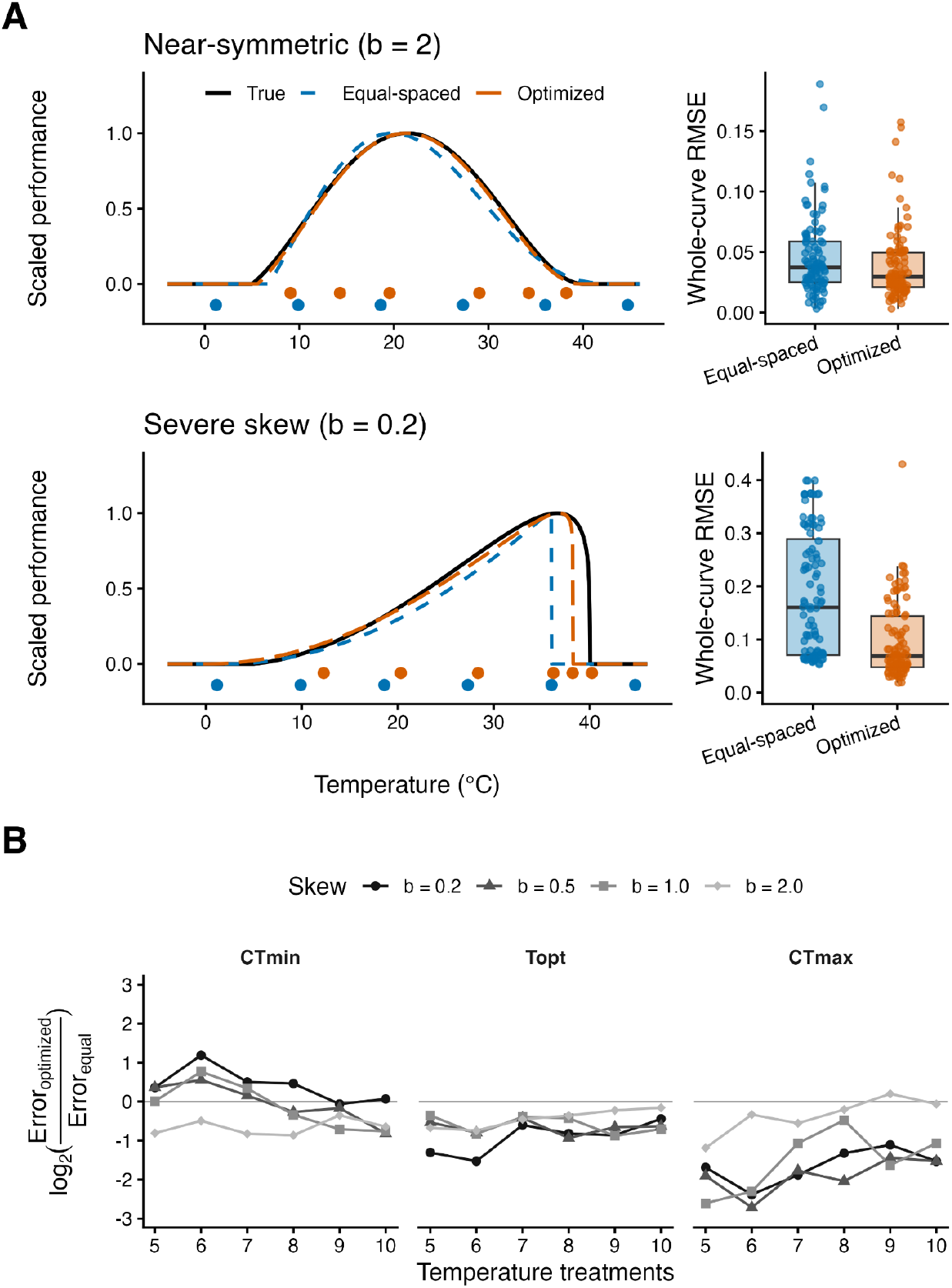
Optimized temperature designs for recovering thermal performance curves with different degrees of skew. A) In each panel, the solid black line shows the true curve, the orange dashed line shows the fitted curve from the optimized design, and the blue dashed line shows the fitted curve from an equal-spaced design. Panels differ only in the skew parameter (b = 2 on top and 0.2 on bottom). Points below the x-axis indicate the temperatures selected under each design. B) The impact of skew on the relative error in estimating *CT*_*min*_, *T*_*opt*_, and *CT*_*max*_ between optimized and equal designs across increasing numbers of temperature treatments. In all panels, unless otherwise specified, curves were generated with *CT*_*min*_ = 5, *CT*_*max*_ = 40, a = 1, six temperature treatments, five replicate observations per temperature, and observation error *obs_cv* = 0.2.

We next examined how experimental performance changed under resource constraints by varying the number of distinct temperature treatments and the number of replicates within these treatments. As expected, increasing experimental effort generally reduced error for both optimized and equally spaced designs, particularly for increasing temperature treatments. Across the full range of treatment and replication combinations examined, optimized designs generally produced lower RMSE than equally spaced designs (Figure 3). Sensitivity analyses exploring additional parameter settings, including uncertainty in prior TPC parameters, are presented in Supplement S2.

**Figure 3.**
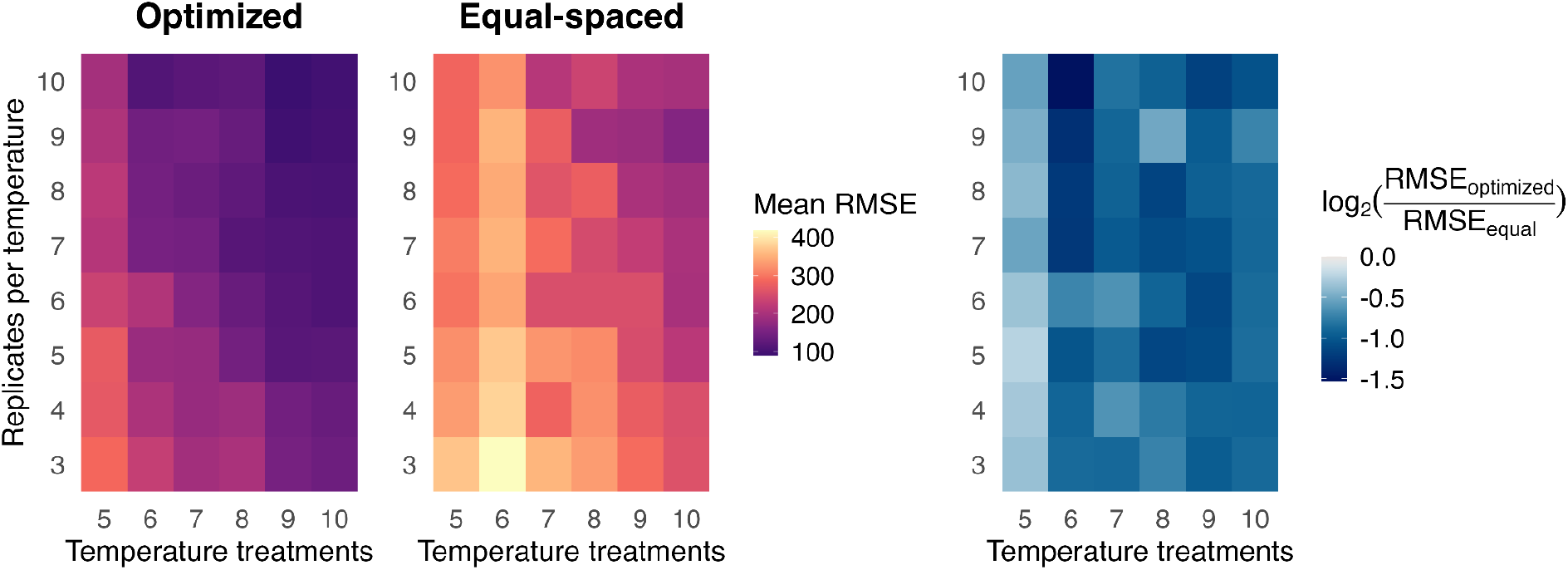
Impact of resource constraints. Heatmap showing error (RMSE) as a function of the number of temperature treatments and replicates for the A) optimized best design, B) equal spaced design, and C) a relative comparison of the two. *CT*_*min*_ *= 5, CT*_*max*_ *= 40, b = 0*.*5, obs_cv = 0*.*2*.

## Discussion

Temperature–trait experiments are often resource intensive, but decisions about where to place temperature treatments are commonly based on convenience or equal spacing rather than on the information those treatments are expected to provide. Our results show that equal spacing is a useful baseline, particularly when TPCs are approximately symmetric or when very few temperature treatments are available, but it is not always the most efficient design. By explicitly accounting for nonlinear curve shape, *TPCdesign* can redistribute experimental effort toward temperatures that are most informative for estimating the full response. This can improve TPC recovery without requiring additional experimental units.

The benefit of optimization depends strongly on TPC shape. Typical ectotherm TPCs are asymmetric, with a relatively gradual rise in performance below the optimum followed by a much steeper decline at high temperatures (Buckley et al., 2021). Under this geometry, successive temperatures along the gradual lower-temperature portion of the curve can provide relatively redundant information, whereas a small change in temperature near the upper shoulder or thermal limit may correspond to a large change in performance. Accordingly, optimized designs tend to place more observations around critical changes in slope and at temperatures that anchor the limits of the response. The advantage over equal spacing therefore becomes greater as asymmetry increases. This principle is consistent with more general work on sampling ecological gradients, which shows that informative locations depend on the underlying response shape and that sampling critical portions of nonlinear responses can improve prediction when prior information about that shape is available (Schweiger et al., 2016; Schweiger et al., 2026).

Resource constraints also determine how experimental effort is divided between the number of temperatures sampled and replication within temperatures. For nonlinear responses, these forms of effort provide different kinds of information. Replication improves precision at temperatures that have already been sampled, but additional temperature treatments are needed to resolve curvature, optima, and thermal limits. As a result, increasing replication cannot fully compensate for a design with too few distinct temperatures, a general problem in experiments along continuous environmental gradients (Kreyling et al., 2018). Our simulations show that optimized temperature placement can reduce estimation error across a broad range of treatment numbers and replication levels, helping make better use of limited experimental effort. Multiple temperature combinations may also provide nearly equivalent expected information, offering flexibility for practical constraints. This is particularly relevant for TPC studies because many existing datasets are sparsely resolved. In the database compiled by Dell et al. (2013), thermal responses were often measured at only a handful of temperatures and frequently covered only one side of the performance curve, limiting the information available to distinguish among nonlinear models and estimate key thermal parameters (Low-Décarie et al., 2017). Kontopoulos et al. (2024) similarly treated datasets with only four distinct temperatures as low resolution and emphasized the need for more temperature levels than fitted model parameters. Although substantially more resolved designs are possible, including experiments spanning 12 temperatures in *Drosophila* (Klepsatel et al., 2023), such intensive sampling remains uncommon.

Another major advantage of this package is the ability to incorporate prior information about thermal performance parameters and specify the uncertainty surrounding those estimates. Optimized designs were generally robust to biased estimates of *CT*min, *CT*max, and *b*, outperforming equally spaced designs across a range of misspecified thermal limits and curve shapes (Supplement S2). When the true curve is skewed, the advantage was weakest when the assumed thermal range was shifted toward warmer temperatures, indicating that the direction of bias matters (Figure S1). Allowing moderate uncertainty around prior estimates generally supported better performance, suggesting that users can benefit from prior knowledge without treating those estimates as precise.

The current implementation also has limits that suggest useful extensions. *TPCdesign* focuses on the Briere-2 model and traits whose performance approaches zero outside lower and upper thermal limits. This assumption will not suit every response. For example, net population growth in microorganisms may become negative beyond temperatures that maximize growth, making observations outside nominal zero-performance limits biologically informative (Luhring & DeLong, 2016; Robey et al., 2026). Different functional forms may also prioritize different parts of the thermal gradient. Recent comparisons have shown substantial variation in how alternative models fit empirical TPCs (Low-Décarie et al., 2017; Padfield et al., 2021; Kontopoulos et al., 2024), suggesting that extending optimal-design methods to additional TPC functions and model-averaged design criteria would be valuable. Additionally, the current method is built on optimizing the full curve fit, but future extensions could instead optimize fitting specific parameters like Topt, CTmax or a specific temperature window.

*TPCdesign* provides a reproducible way to use existing biological knowledge to make informed decisions about experimental designs. By planning temperature placement alongside replication and other resource constraints, researchers can obtain more informative TPCs from the same experimental effort and strengthen the physiological parameters on which ecological forecasts increasingly depend.

## Author contributions

K.L. and L.A.N. conceived the study. L.A.N. developed the *TPCdesign* software and implemented the optimization framework. K.L. and L.A.N. developed the simulation analyses and visualization. K.L. led manuscript preparation. All authors contributed critically to manuscript development and revisions.

## Supplement

### S1. Baseline uncertainty in priors from within genus comparisons

To estimate how accurately users could approximate Brière-2 parameters from related species, we performed a leave-one-species-out validation on the dataset from Buckley et al. (2021). For each focal species in a genus or family with at least three species with valid Tmin, Tmax, and derived b, we temporarily removed that species from the group. We then predicted its Tmin, Tmax, and b as the median value of the remaining species in the same genus or family. Prediction error was calculated as the absolute difference between the observed species value and this related-species median. We summarized these errors across all eligible species using the median absolute error. We included 83 genus-level species and 204 family-level species. Eligible groups spanned multiple organism types, including algal/phytoplankton, insects and other arthropods, fishes, plants, and lizards.

### S2. Exploration of prior uncertainty

We evaluated the sensitivity of optimized temperature designs to uncertainty and bias in prior estimates of thermal performance parameters. The true curve was defined by *CT*min = 5°C, *CT*max = 40°C, and *b* = 0.5. Prior estimates varied across all combinations of offsets of −2.5, 0, or +2.5°C for *CT*min; −1.5, 0, or +1.5°C for *CT*max; and −0.26, 0, or +0.26 for log(*b*). We considered five prior-uncertainty multipliers (0.5, 1, 1.5, 2, and 3), for a total of 135 conditions, with 100 simulated fits per design method in each condition. Performance was summarized using mean whole-curve RMSE.

**Figure S1.**
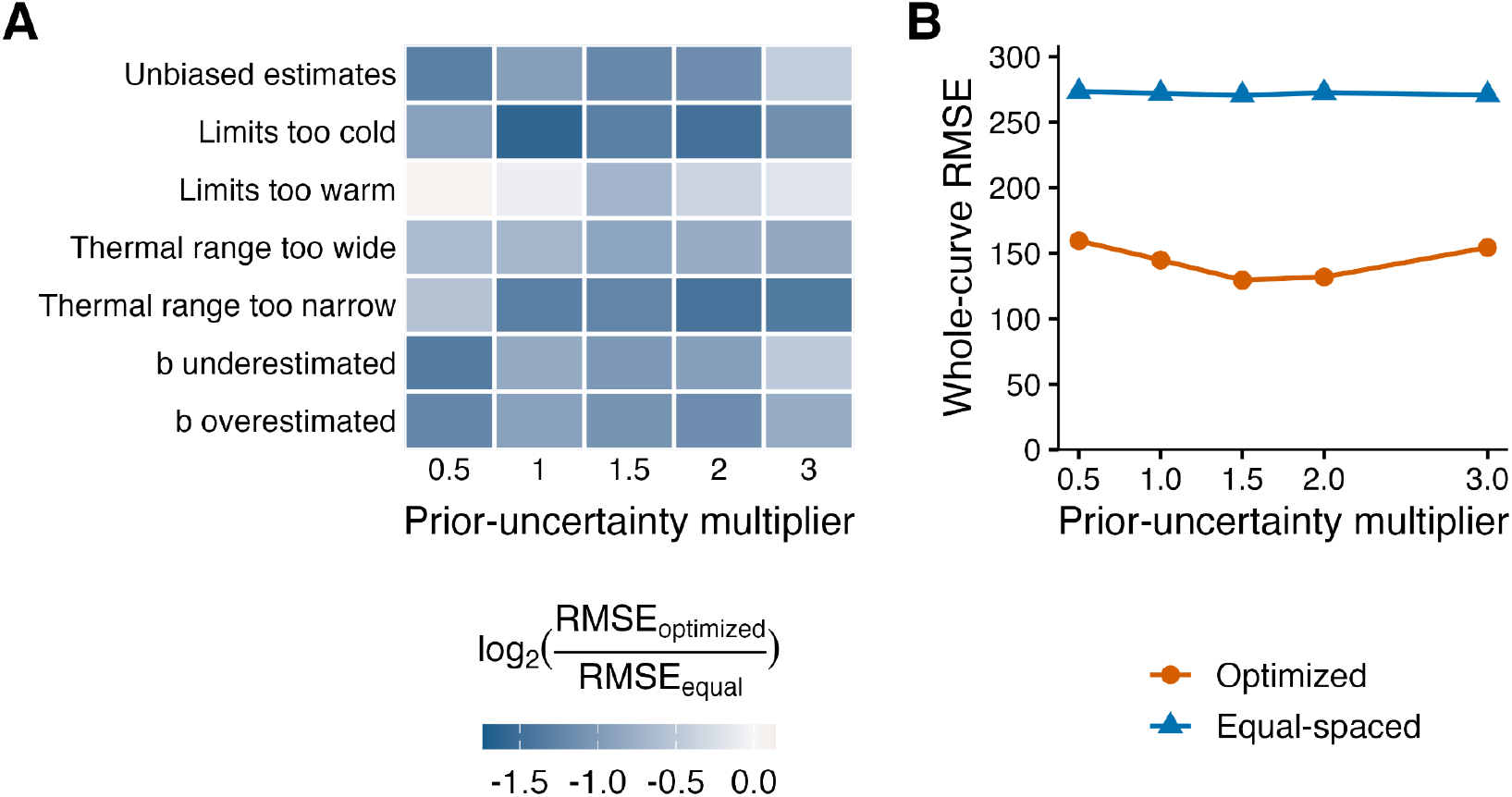
Sensitivity to prior bias and uncertainty. (A) Ratio of RMSE for optimized versus equally spaced designs under seven representative scenarios: unbiased estimates, thermal limits shifted toward colder or warmer temperatures, an assumed thermal range that was too wide or too narrow, and underestimated or overestimated *b*. Darker blue indicates a greater advantage for optimized designs; a ratio of 1 indicates equal performance. (B) Mean whole-curve RMSE for each design method, pooled across all 27 offset combinations at each uncertainty multiplier. Optimized designs generally retained an advantage despite biased prior estimates, with the weakest advantage when the assumed thermal range was shifted toward warmer temperatures. Intermediate uncertainty settings produced the lowest pooled mean RMSE.

## Notes

### Competing Interest Statement

The authors have declared no competing interest.

## References

Angilletta, M. (2009). Thermal Adaptation: A Theoretical and Empirical Synthesis. Oxford University Press, Oxford.

Bengtsson, H. (2021). A unifying framework for parallel and distributed processing in R using futures. The R Journal, 13, 208–227. 10.32614/RJ-2021-048

Bernhardt, J.R., Sunday, J.M., Thompson, P.L. & O’Connor, M.I. (2018). Nonlinear averaging of thermal experience predicts population growth rates in a thermally variable environment. Proceedings of the Royal Society B: Biological Sciences, 285, 20181076. 10.1098/rspb.2018.1076

Briere, J.F., Pracros, P., Le Roux, A.Y. & Pierre, J.S. (1999). A novel rate model of temperature-dependent development for arthropods. Environmental Entomology, 28, 22–29. 10.1093/ee/28.1.22

Buckley, L.B., Huey, R.B. & Kingsolver, J.G. (2022). Asymmetry of thermal sensitivity and the thermal risk of climate change. Global Ecology and Biogeography, 31, 2231–2244. 10.1111/geb.13570

Carnell, R. (2026). lhs: Latin hypercube samples. R package version 1.2.1. 10.32614/CRAN.package.lhs

Chaloner, K. & Verdinelli, I. (1995). Bayesian experimental design: A review. Statistical Science, 10, 273–304. 10.1214/ss/1177009939

Dell, A.I., Pawar, S. & Savage, V.M. (2013). The thermal dependence of biological traits. Ecology, 94, 1205–1206. 10.1890/12-2060.1

Eddelbuettel, D. & François, R. (2011). Rcpp: Seamless R and C++ integration. Journal of Statistical Software, 40, 1–18. 10.18637/jss.v040.i08

Eddelbuettel, D. & Sanderson, C. (2014). RcppArmadillo: Accelerating R with high-performance C++ linear algebra. Computational Statistics and Data Analysis, 71, 1054–1063. 10.1016/j.csda.2013.02.005

Huey, R.B. & Kingsolver, J.G. (1989). Evolution of thermal sensitivity of ectotherm performance. Trends in Ecology & Evolution, 4, 131–135. 10.1016/0169-5347(89)90211-5

Kontopoulos, D.G., Sentis, A., Daufresne, M., Glazman, N., Dell, A.I. & Pawar, S. (2024). No universal mathematical model for thermal performance curves across traits and taxonomic groups. Nature Communications, 15, 8855. 10.1038/s41467-024-53046-2

Krenek, S., Berendonk, T.U. & Petzoldt, T. (2011). Thermal performance curves of Paramecium caudatum: A model selection approach. European Journal of Protistology, 47, 124–137. 10.1016/j.ejop.2010.12.001

Kreyling, J., Schweiger, A.H., Bahn, M., Ineson, P., Migliavacca, M., Morel-Journel, T., Christiansen, J.R., Schtickzelle, N. & Larsen, K.S. (2018). To replicate, or not to replicate – that is the question: How to tackle nonlinear responses in ecological experiments. Ecology Letters, 21, 1629–1638. 10.1111/ele.13134

Low-Décarie, E., Boatman, T.G., Bennett, N., Passfield, W., Gavalás-Olea, A., Siegel, P. & Geider, R.J. (2017). Predictions of response to temperature are contingent on model choice and data quality. Ecology and Evolution, 7, 10467–10481. 10.1002/ece3.3576

Luhring, T.M. & DeLong, J.P. (2016). Predation changes the shape of thermal performance curves for population growth rate. Current Zoology, 62, 501–505. 10.1093/cz/zow045

MacLean, H.J., Sørensen, J.G., Kristensen, T.N., Loeschcke, V., Beedholm, K., Kellermann, V. & Overgaard, J. (2019). Evolution and plasticity of thermal performance: An analysis of variation in thermal tolerance and fitness in 22 Drosophila species. Philosophical Transactions of the Royal Society B: Biological Sciences, 374, 20180548. 10.1098/rstb.2018.0548

McKay, M.D., Beckman, R.J. & Conover, W.J. (1979). A comparison of three methods for selecting values of input variables in the analysis of output from a computer code. Technometrics, 21, 239–245. 10.1080/00401706.1979.10489755

Molnár, P.K., Sckrabulis, J.P., Altman, K.A. & Raffel, T.R. (2017). Thermal performance curves and the metabolic theory of ecology: A practical guide to models and experiments for parasitologists. Journal of Parasitology, 103, 423–439. 10.1645/16-148

Overgaard, J., Kearney, M.R. & Hoffmann, A.A. (2014). Sensitivity to thermal extremes in Australian Drosophilaimplies similar impacts of climate change on the distribution of widespread and tropical species. Global Change Biology, 20, 1738–1750. 10.1111/gcb.12521

Overstall, A.M. & Woods, D.C. (2017). Bayesian design of experiments using approximate coordinate exchange. Technometrics, 59, 458–470. 10.1080/00401706.2016.1251495

Overstall, A.M., Woods, D.C. & Adamou, M. (2020). acebayes: An R package for Bayesian optimal design of experiments via approximate coordinate exchange. Journal of Statistical Software, 95, 1–33. 10.18637/jss.v095.i13

Paaijmans, K.P., Heinig, R.L., Seliga, R.A., Blanford, J.I., Blanford, S., Murdock, C.C. & Thomas, M.B. (2013). Temperature variation makes ectotherms more sensitive to climate change. Global Change Biology, 19, 2373–2380. 10.1111/gcb.12240

Padfield, D., O’Sullivan, H. & Pawar, S. (2021). rTPC and nls.multstart: A new pipeline to fit thermal performance curves in R. Methods in Ecology and Evolution, 12, 1138–1143. 10.1111/2041-210X.13585

Quinn, B.K. (2017). A critical review of the use and performance of different function types for modeling temperature-dependent development of arthropod larvae. Journal of Thermal Biology, 63, 65–77. 10.1016/j.jtherbio.2016.11.013

Rebaudo, F., & Rabhi, V.B. (2018). Modeling temperature-dependent development rate and phenology in insects: Review of major developments, challenges, and future directions. Entomologia Experimentalis et Applicata, 166, 607–617. 10.1111/eea.12693

Régnière, J., Powell, J., Bentz, B. & Nealis, V. (2012). Effects of temperature on development, survival and reproduction of insects: Experimental design, data analysis and modeling. Journal of Insect Physiology, 58, 634–647. 10.1016/j.jinsphys.2012.01.010

Robey, A.J., Kummel, M.T. & Vasseur, D.A. (2026). Temporal autocorrelation increases temperature-driven extinction risk by clustering stressful conditions. Ecology Letters, 29, e70441. 10.1111/ele.70441

Schweiger, A.H., Garthen, A., Bahn, M., Chalcraft, D., Schtickzelle, N., Larsen, K.S. & Kreyling, J. (2026). How to optimally allocate sampling effort in experimental ecology. Scientific Reports, 16, 6503. 10.1038/s41598-026-38541-4

Schweiger, A.H., Irl, S.D., Steinbauer, M.J., Dengler, J. & Beierkuhnlein, C. (2016). Optimizing sampling approaches along ecological gradients. Methods in Ecology and Evolution, 7, 463–471. 10.1111/2041-210X.12495

Sinclair, B.J., Marshall, K.E., Sewell, M.A., Levesque, D.L., Willett, C.S., Slotsbo, S., Dong, Y., Harley, C.D.G., Marshall, D.J., Helmuth, B.S. & Huey, R.B. (2016). Can we predict ectotherm responses to climate change using thermal performance curves and body temperatures? Ecology Letters, 19, 1372–1385. 10.1111/ele.12686

Sorek, S., Smith, J.W., Jr., Huxley, P.J. & Johnson, L.R. (2025). bayesTPC: Bayesian inference for thermal performance curves in R. Methods in Ecology and Evolution, 16, 687–697. 10.1111/2041-210X.70004

Vasseur, D.A., DeLong, J.P., Gilbert, B., Greig, H.S., Harley, C.D.G., McCann, K.S., Savage, V., Tunney, T.D. & O’Connor, M.I. (2014). Increased temperature variation poses a greater risk to species than climate warming. Proceedings of the Royal Society B: Biological Sciences, 281, 20132612. 10.1098/rspb.2013.2612

